# Integrated analyses of the single-cell ATAC-seq and RNA-seq reveal the epigenetic landscape of human ovarian aging

**DOI:** 10.1101/2021.11.07.467593

**Authors:** Yunzhao Xu, Jinling Chen, Shuting Gu, Yuanlin Liu, Huihua Ni, Yan Huang, Sainan Gao, Nan Sheng, Xiaojing Zhang, Xin He, Songlin Zhou, Wenliang Ge

## Abstract

Studying the molecular mechanisms of ovarian aging is crucial for understanding the age-related fertility issues in females. Recently, a single-cell transcriptomic roadmap of ovarian aging based on non-human primates revealed the molecular signatures of the oocytes at different developmental stages. Herein, we present the first epigenetic landscape of human ovarian aging, through an integrated analysis of the single-cell assay for transposase-accessible chromatin using sequencing (scATAC-seq) and single-cell RNA-seq. We depicted the transcriptional profiles and chromatin accessibility of the ovarian tissues isolated from old (n=4) and young (n=2) donors. The unsupervised clustering of data revealed seven distinct cell populations in the ovarian tissues and six subtypes of oocytes, which could be distinguished by age difference. Further analysis of the scATAC-seq data from the young and old oocytes revealed that the interaction between the Notch signaling pathway and AP-1 family transcription factors may crucially determine oocyte aging. Finally, a machine-learning algorithm was applied to calculate the optimal model based on the single-cell dataset for predicting oocyte aging, which exhibited excellent accuracy with a cross-validated area under the receiver operating characteristics score of 0.99. In summary, this study provides a comprehensive understanding of human ovarian aging at both the transcriptomic and epigenetic levels, based on an integrated analysis of large-scale single-cell datasets. We believe our results will shed light on the discovery of potential therapeutic targets or diagnostic markers for age-related ovarian disorders.

## Itroduction

The ovary is the chief female reproductive organ that not only stores and provides oocytes but also supplies essential sex hormones, fundamentally determining the reproductive lifespan and maintaining the endocrine homeostasis in females^1^. However, women are born with a finite number of follicles^2^. As the number of follicles decreases with age, female fertility declines and eventually leads to menopause as a physiological consequence of ovarian aging^3^. Also, the specific reason for the cessation of ovulation in females after 35 years of age remains unknown^4^. Ovarian aging is also closely related to a wide spectrum of diseases, including ovarian cancer, breast cancer, cardiovascular disease, and type 2 diabetes mellitus^5,6^. This necessitates the elucidation of the mechanism of ovarian aging and identification of the possible target to delay the process.

Ovary possesses a complex structure with a heterogeneous population of cells at progressive developmental stages^7^. The follicle is the main functional unit of the ovary^8^. Each follicle consists of a partially differentiated oocyte, enclosed within increasing layers of somatic granulosa cells and theca-interstitial cells that support the oocyte during maturation^1^. Other components of the ovary include the stromal cells, smooth muscle cells, endothelial cells, and various immune cells^9,10^. Therefore, unraveling the cellular landscape of the ovary is indispensable for a better understanding of cell-type-specific dynamics during ovarian aging.

Advances in single-cell RNA sequencing (scRNA-seq) technology enable the cellular landscape to be mapped at the single-cell level and the cell trajectory to be inferred for examining the dynamic changes in the cell populations^11^. Previous studies performed the scRNA-seq on the ovaries of young and old cynomolgus monkeys^7^. Researchers have depicted the transcriptional profile of four oocyte subtypes and demonstrated that oxidative damage and subsequent responses are essential for the aging-related changes in early-stage oocytes and granulosa cells^7^. Another study focused on the growth and regression of the follicular populations in human adult ovaries^10^. Various cell types have been identified, and a significant association was found between the complement system and follicular remodeling^10^. However, the epigenetic landscape of ovarian aging in humans is unclear.

In addition to the transcriptional regulation, epigenetic reprogramming also takes place in the postnatal oocyte growth and ovarian aging. Previous studies reported that persistence of methylation at CpG islands affected the expression of genes related to the preimplantation development in oocytes^12,13^. Nevertheless, the chromatin landscape and its dynamics during the ovarian aging are still unexplored. Single cell assay for transposase-accessible chromatin using sequencing (scATAC-seq) is an exquisite tool for sketching the chromatin accessibility profile at single-cell resolution^14^. Combined with the scRNA-seq, the comprehensive analysis through scATAC-seq is promising for unveiling the epigenetic mechanism of ovarian aging.

## Results

### Single-cell sequencing of over 49,000 cells from the ovarian tissues of six patients

To study the transcriptomic heterogeneity and compare the old and young ovarian tissues based on the cell types, the tissues were collected from six patients (**Materials and Methods, Supplementary Figure 1**) and the single-cell sequencing was performed using the 10x Genomics platform. After the quality control and batch integration, 49,651 cells were clustered into ten initial clusters (**Figure 1A**). The marker genes from the literature were used to annotate these clusters, including CDKN1C for the stromal cells, TAGLN and SOD3 for the oocytes, SPARC and TIMP1 for the smooth muscle cells, INHA for the granulosa cells, CDH5 and VWF for the endothelial cells, CD14 and CD74 for the macrophages, IL32 and CCL5 for the natural killer (NK) cells (**Figure 1B**). The clusters were then integrated and renamed based on the marker genes and clustered cells into seven cell types (**Figure 1C**). The top markers of each cell type were calculated and showed clear patterns for each cell type (**Figure 1D**). To further validate the annotations of the clusters, the functional enrichment analyses were performed for the significantly highly expressed genes for each cluster (*p*-value < 0.05, fold-change > 1.2) (**Supplementary Figure 2**). As expected, the enriched Gene Ontology (GO) terms revealed critical functions of the corresponding cells. For example, the top enriched GO terms for the macrophages and NK cells were mainly immune-related pathways, such as “regulation of immune response,” “immune response-activating signal transduction” and “activation of immune response.”

**Figure 1.**
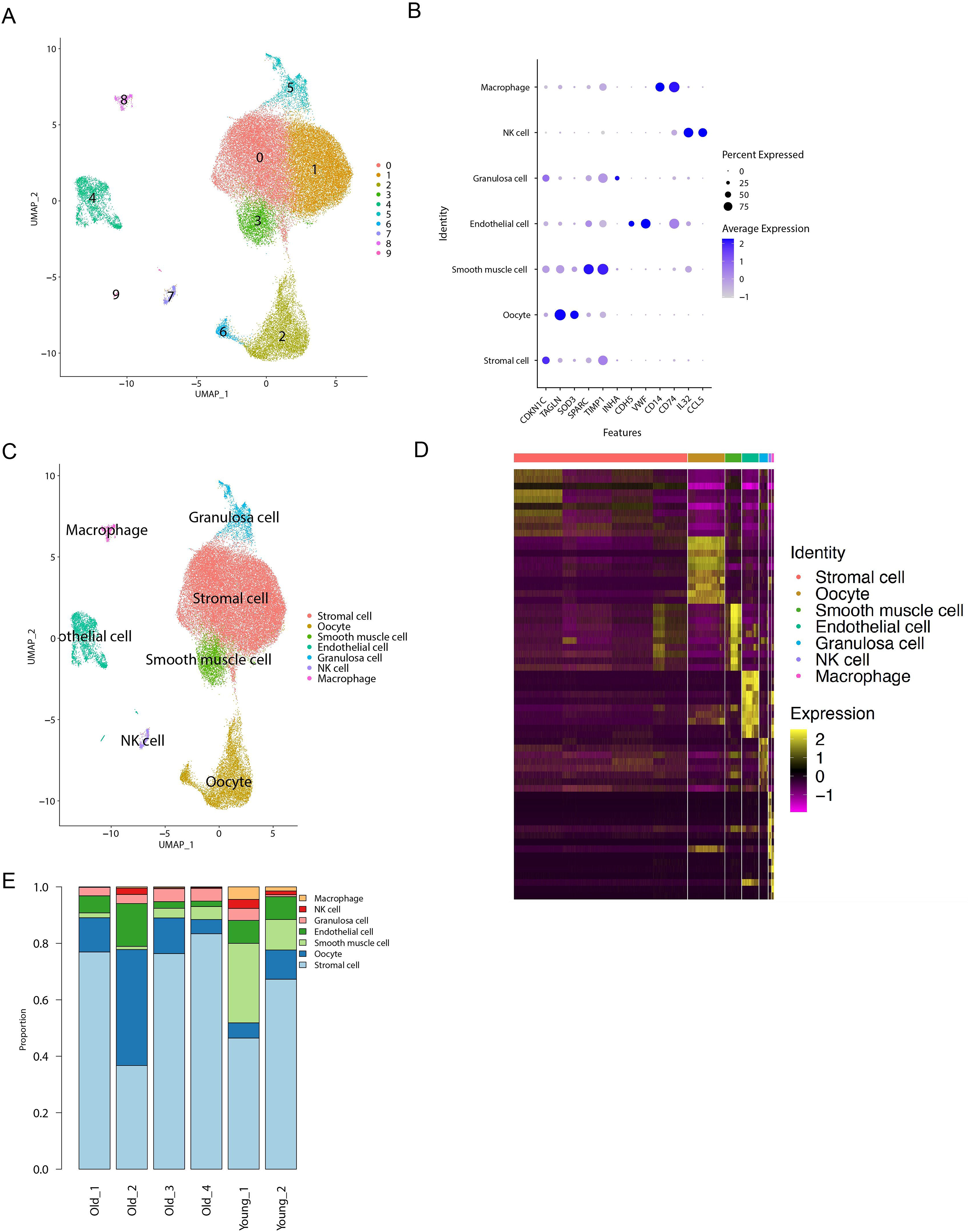
Single-cell clustering and annotation. A. UMAP visualization of initial clustering results. B. Marker genes reported by the literature (Materials and Methods) were used to define clusters. For gene expression, the raw read counts were normalized and transformed. C. UMAP visualization with cell-type annotation results. D. Top 10 marker genes for each cell type visualized in a heatmap. For gene expression, the raw read counts were normalized and transformed, and then scaled. E. Proportions of each cell type in each patient sample.

We further explored the changes in the composition of each cell type between the old and young ovarian samples (**Figure 1E**). It is worth mentioning that the number of macrophages and NK cells was significantly decreased in the old patient samples compared to the young patient samples, while the stromal cells increased in the old patient samples. Aging has been reported to impair many aspects of macrophage functions^15–17^ and has also been identified to have critical impacts on the macrophage-involved immune response.

### Six subtypes of oocytes show distinct gene expression signatures at the sequential development stage

We conducted an unsupervised analysis of oocyte gene expression profiles to explore the dynamic changes in the oocyte transcriptomic profiles and identified six subtypes of oocytes (C0 to C5). Uniform Manifold Approximation and Projection (UMAP) revealed that the six subtypes were distributed along the UMAP1 dimension (**Figure 2A**). To investigate the dynamic changes in gene expression during oocyte development, we calculated a subset of feature genes that exhibited high subtype-to-subtype variations (**Figure 2B**). Notably, we found that most of the C1–C2 oocytes were from young tissues, while the C3–C5 oocytes were from the old. We subsequently examined the gene expression across the six subtypes and identified the top 10 highly expressed genes for each subtype (**Figure 2D**). To validate the functional difference of each subtype, we conducted the GO analysis for the significantly highly expressed genes for each cell type (*p*< 0.05) (**Figure 2D**). Consistent with the previous reports and the above canonical gene expression patterns, the GO analysis revealed the biological processes that were enriched for each stage of follicular development. Notably, the “protein targeting to endoplasmic reticulum” and “ribosome biogenesis” were enriched for subtype C0 with high expression of ribosomal protein-related genes (representative genes *RPS8*, *RPS9*, and *RPS14*). We also identified that “oxidative phosphorylation” (representative genes *ATP5ME* and *COX5B*) and “ATP synthesis coupled electron transport” (representative genes *NDUFA4* and *COX6C*) were enriched for the subtype C4. The variety of enriched GO terms revealed the functional heterogeneity of these subtypes during folliculogenesis. As shown in **Figure 2B**, we identified a subset of feature genes that exhibited high subtype-to-subtype variations. We then examined the expression trends of these feature genes and identified the genes that gradually increased or decreased with oocyte development. It is worth mentioning that we found that two critical oocyte-related genes, *TIMP1* and *TIMP2*, had significant expression trends in correlation with oocyte development (**Figure 2E**). Previous studies have reported that TIMP1 and TIMP2 are associated with oocyte maturity and quality and are necessary for normal follicular development ^18,19^. In our results, TIMP1 and TIMP2 displayed a gradually decreasing expression along with oocyte development.

**Figure 2.**
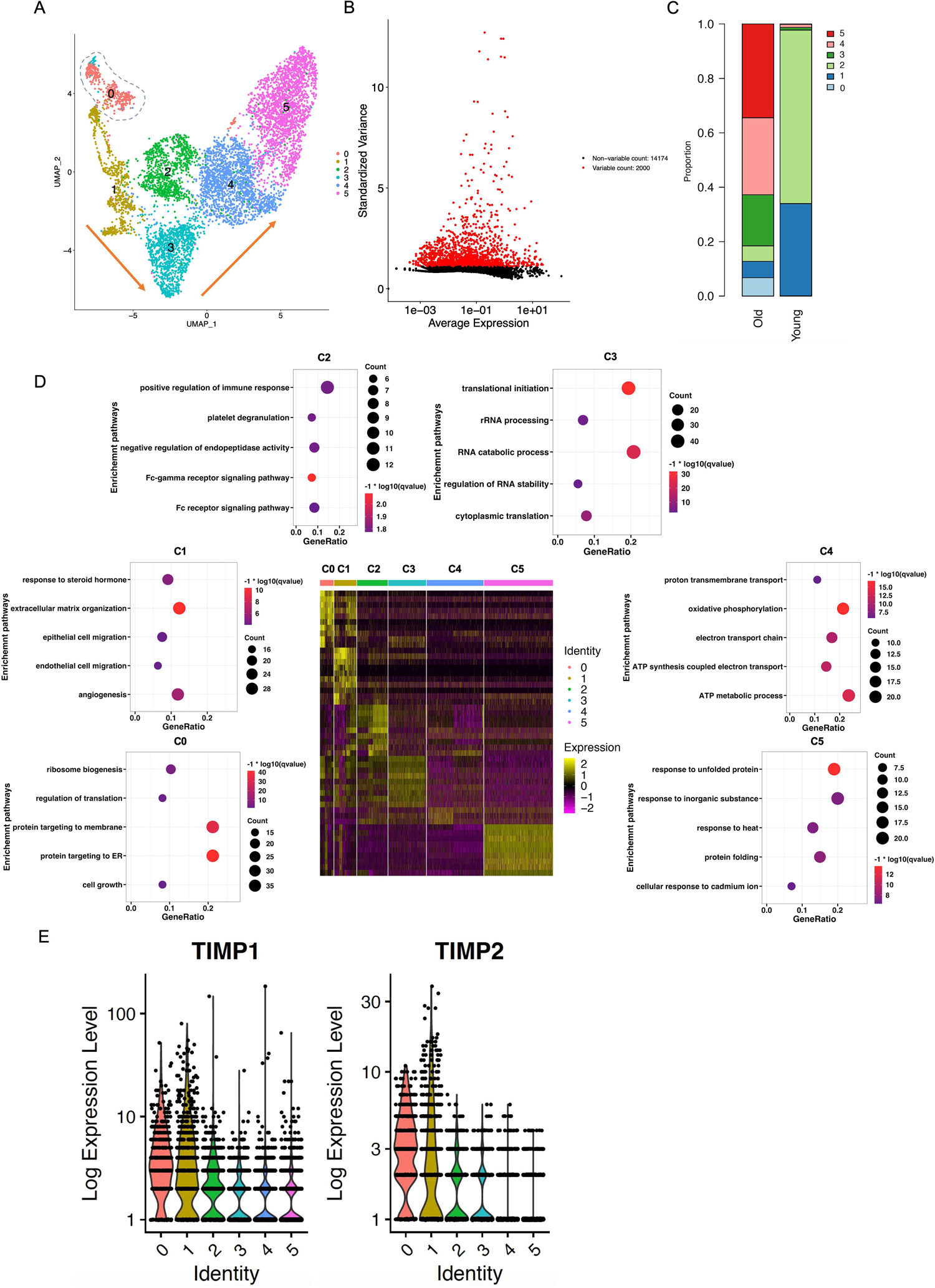
A single-cell cluster of oocyte cells. A. UMAP of six subclusters of the oocyte. B. Subset of genes with high variation in subtype-to-subtype transition. C. Proportion of six subclusters of oocyte in the young and old patient samples. Subcluster C0 = identity 0, C1 = identity 1 … C5 =identity 5, respectively. C. Heatmap of six oocyte subtypes and their corresponding GO enrichment results. D. Violin plots of TIMP1 and TIMP2.

### The scATAC-seq profiling of human ovaries identified different cell types and gene expression signatures

To profile the genome-wide chromatin accessibility of the cells from the old and young ovarian tissues, a scATAC-seq approach was applied to investigate the differences between the old and young ovarian samples. We utilized “ArchR,” a state-of-the-art scATAC-seq analysis R package, for quality control, doublet removal, cell clustering, and annotation, peak calling, differential chromatin accessibility analysis, and footprinting analysis were performed^20^. Cell barcodes with more than 4,000 total unique fragments and a transcription start site enrichment score of more than 8 were selected for downstream analysis (**Supplementary Figures3A and B**). After quality control and removal of doublets, a total of 38,312 single cells were obtained from four old and two young patients’ samples (**Supplementary Figures3C, 4, and 5**). For the downstream cell clustering and visualization, the graph-based Louvain algorithm and UMAP were implemented in the ‘Seurat’ R package. Based on chromatin accessibility profiling, 38,312 cells were represented by a total of 16 clusters (**Figure 3A**). To accurately annotate the clusters, the aforementioned scRNA-seq annotation information was integrated into our scATAC-seq analysis. Finally, seven distinct cell clusters representing the cell types were discovered by scRNA-seq data, including endothelial cells, granulosa cells, macrophages, NK cells, oocytes, smooth muscle cells, and stromal cells (**Figure 3B**). By using ArchR’s default peak caller MACS2 with recommended settings, a total of 258,273 peaks were identified in the old and young ovarian samples. After identification of the robust peak sets for each cell type, we further predicted the type of transcription factors (TFs) that may mediate the binding events creating accessible chromatin sites. The motif enrichment results revealed distinct sets of TFs for cell types, indicating the critical functions of these TFs in regulating the cells (**Figure 3C**). The footprinting analysis also revealed the specific upregulated binding of the TFs for each cell type (**Figure 3D**). For example, the RUNX1 TF has been reported to be associated with the NK cell clonal expansion and memory formation. In addition, STAT6 is a key TF in macrophage M2 polarization^21,22^. Our results were supported by the results of previous studies and reveal the novel functional TFs for each cell type during ovarian aging.

**Figure 3.**
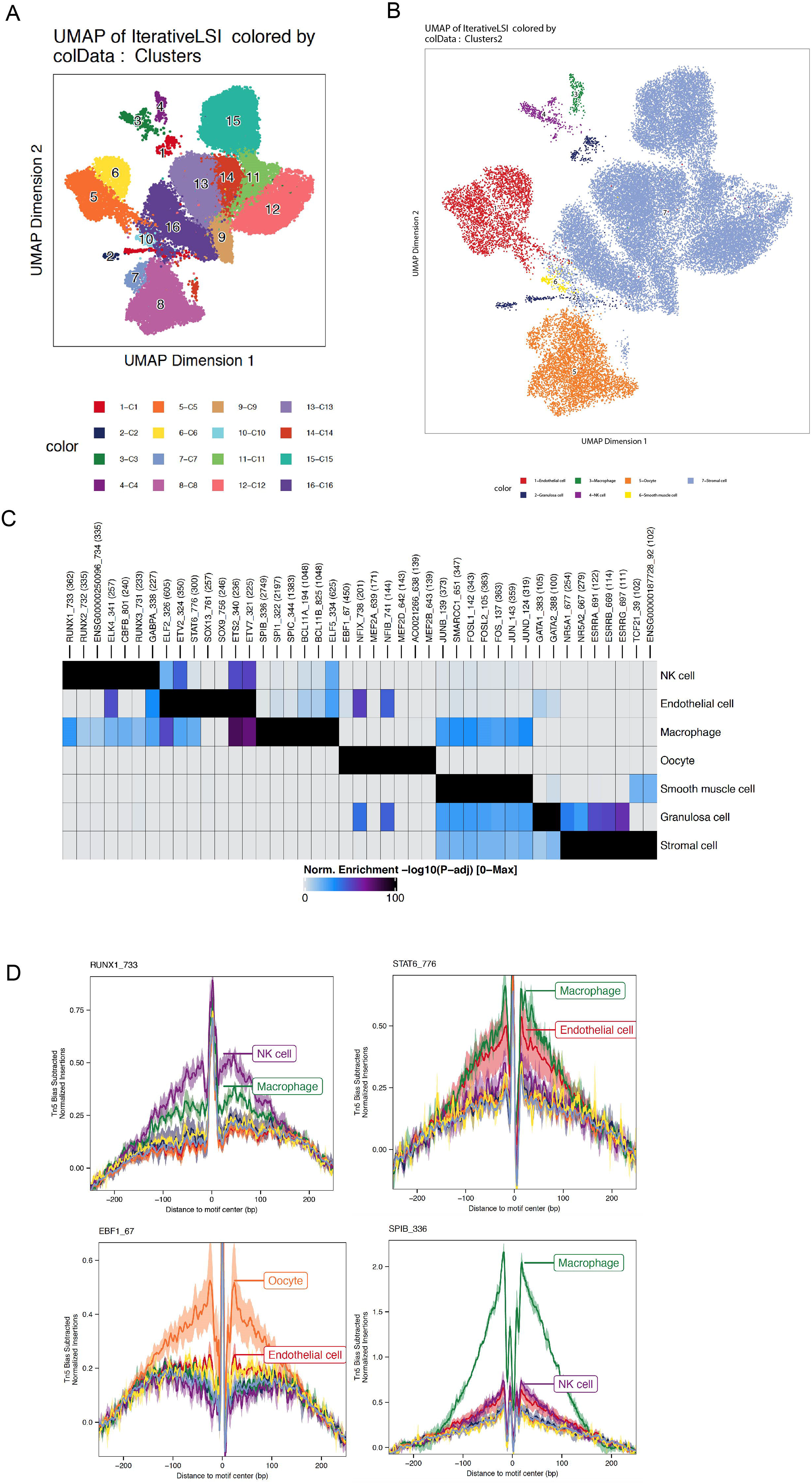
scATAC analysis of ovarian tissues. A. UMAP visualization of initial clustering results. B. UMAP visualization of annotated clustering results. C. Enriched transcription factor motifs for each cell type. D. Footprinting plot of representative transcription factors for each cell type.

### The integrated scRNA-seq and scATAC-seq analyses demonstrate the functional roles of the AP-1 TF family and Notch signaling pathway in ovarian aging

To investigate the molecular mechanisms of oocyte aging, the chromatin accessibility of oocyte cells from old and young patients was further compared. The MACS2 with recommended settings was applied to identify the peaks for the old and young oocyte cells and predict the TFs that mediate the binding events creating accessible chromatin sites (**Figure 4A**). The motif enrichment results revealed distinct sets of TFs in the old and young oocytes, which implied the distinct functional roles of TFs in regulating oocyte aging (**Figure 4B**). Notably, we found that multiple critical AP-1 TF family members, such as FOS, JUNB, FOSL1, and FOSL2, were highly enriched in the old oocyte cells compared to the young oocytes. We further performed the functional enrichment analysis for the differentially enriched TFs between the old and young oocytes. It is worth mentioning that several critical aging pathways, including “AP-1 TF network,” “tumor necrosis factor signaling pathway” and “Aging” terms, were enriched for significantly upregulated TFs in the old oocyte cells (**Figure 4C**). The aging term was enriched with several critical upregulated genes, including *FOS*, *CREB1*, *JUN*, *TP63*, and *PAX2*. For the enriched pathways for upregulated TFs in young oocyte cells, the Notch signaling pathway was notably one of the most significantly enriched terms (**Figure 4D**). AP-1 was reported to negatively regulate the Notch signaling pathway ^23^, which is consistent with our findings, indicating an interaction between these two pathways. In our results, the AP-1 TF family activity was significantly upregulated in the old oocyte cells, whereas the Notch signaling pathway was downregulated. The Notch signaling pathway has been widely reported to be associated with aging and plays important role in age-related diseases^24,25^. Our results support the negative regulation between AP-1 and the Notch signaling pathway and suggest the potential functional roles of the AP-1 TF family in aging. To further validate the upregulation of the AP-1 family in the old oocyte cells, we conducted the footprinting analysis for several critical AP-1 family members and integrated their expression in the scRNA-seq data. We found that FOSL2 and ATF3 showed profoundly upregulated patterns in the old oocyte cells in both the footprinting and scRNA-seq results (**Figure 4E; Supplementary Figure 6**). FOSL2 and ATF3 worked orchestrally in the AP-1 complex. In order to validate this crucial finding, we performed immunostaining assays on both young and old ovarian tissues. Interestingly, the positive signal of AP-1 complex (c-fos and c-jun) can only be found in old oocyte cells, but not in the young oocyte cells (**Figure 4F**). Therefore, together with the experimental validation results, our integrated analysis of the scRNA-seq and scATAC-seq indicated consistent patterns of upregulation of AP-1 signal in oocyte aging, thus indicating the potential role of AP-1 regulation in ovarian aging.

**Figure 4.**
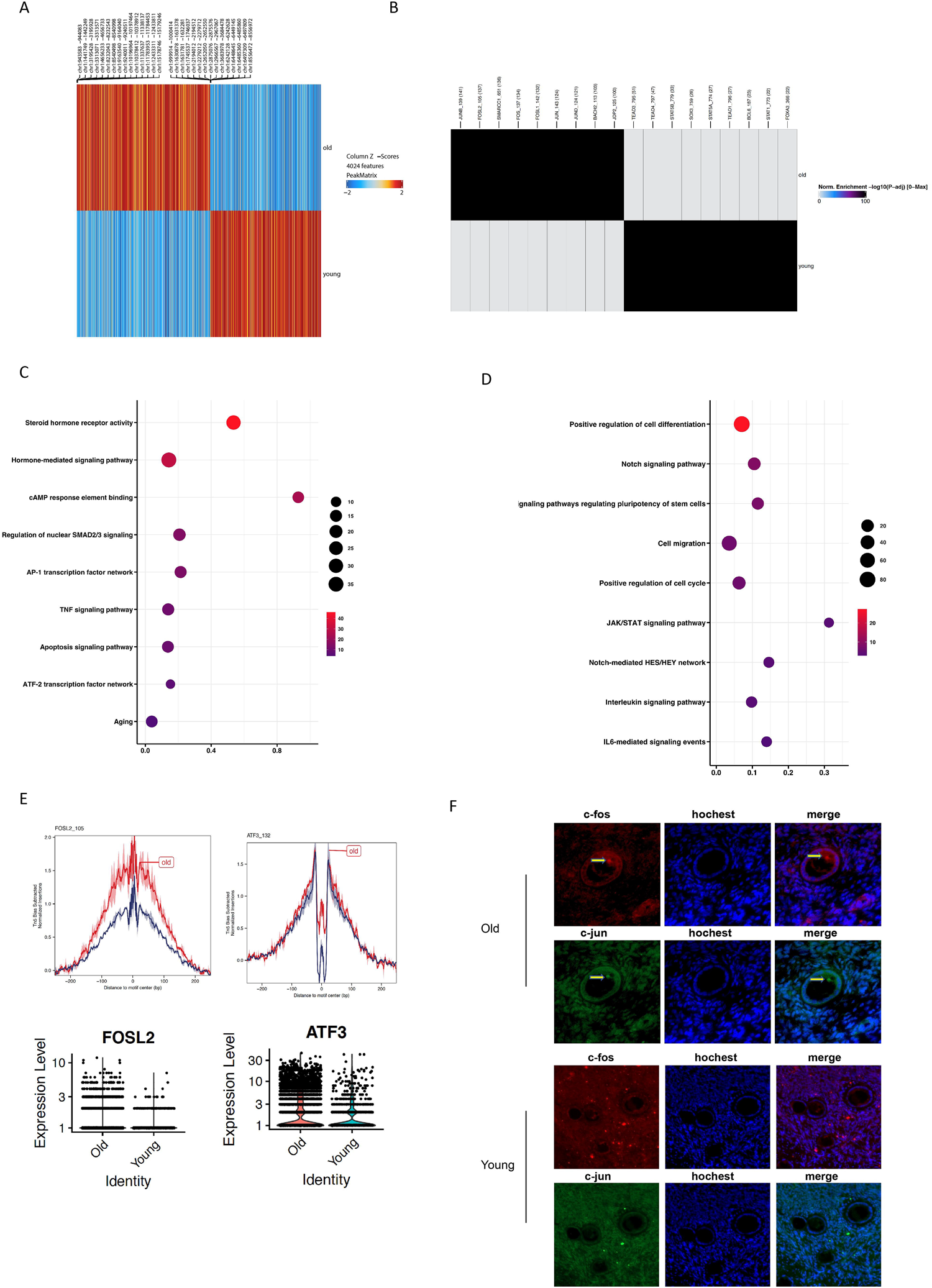
scATAC analysis of the old and young oocyte cells. A. Heatmap of enriched motifs of the old and young oocytes. B. Enriched transcription factors of the old and young oocytes. C-D. Functional enrichment of transcription factor motifs identified for the old and young oocyte cells, respectively. E. Footprinting and violin plots of FOSL2 and ATF3.

#### Machine-learning-based prediction of ovarian aging

It is widely established that bulk RNA-seq data have been used to investigate the biological processes of aging. Recently, expression profiles at the single-cell level have been applied to human skin aging and mouse aging in multiple tissues ^26,27^. We found that the gene expression of oocyte cells was significantly different between the young and old ovaries in our scRNA-seq dataset. Next, we applied our scRNA-seq data to predict ovarian aging. A statistical model was then built and optimized to describe the relationship between gene expression and ovarian aging. The mutual information scores were used to rank the genes. The cross-validated AUROC scores for models with different numbers of top-ranked genes are shown in **Figure 5A**. We included the 10 top-ranked genes in the predictive model, as including more genes did not further improve the performance of the model. The performance of the optimized model (see **Materials and Methods**) was characterized using Receiver Operating Characteristic (ROC) analysis (**Figure 5B**). The model with the selected 10-gene signature exhibited excellent accuracy, with a cross-validated AUROC score of 0.99. The other metrics of the model were also commensurate with the AUROC score (**Figure 5C**). The gene signature included those of *MT-ATP6, MT-CO3, MT-CYB, IGFBP7, VIM, RPL3, RPL27A, HIST1H4C, HLA-A*, and *CD63* (**Figure. 5D**). We also performed the Mann–Whitney–Wilcoxon test with Bonferroni correction and found that all of these genes were significantly differentially expressed between the two groups with *p*-values smaller than 10^-4^. These findings suggest excellent predictability of ovarian aging using the 10-gene signature expression of the oocyte cells.

**Figure 5.**
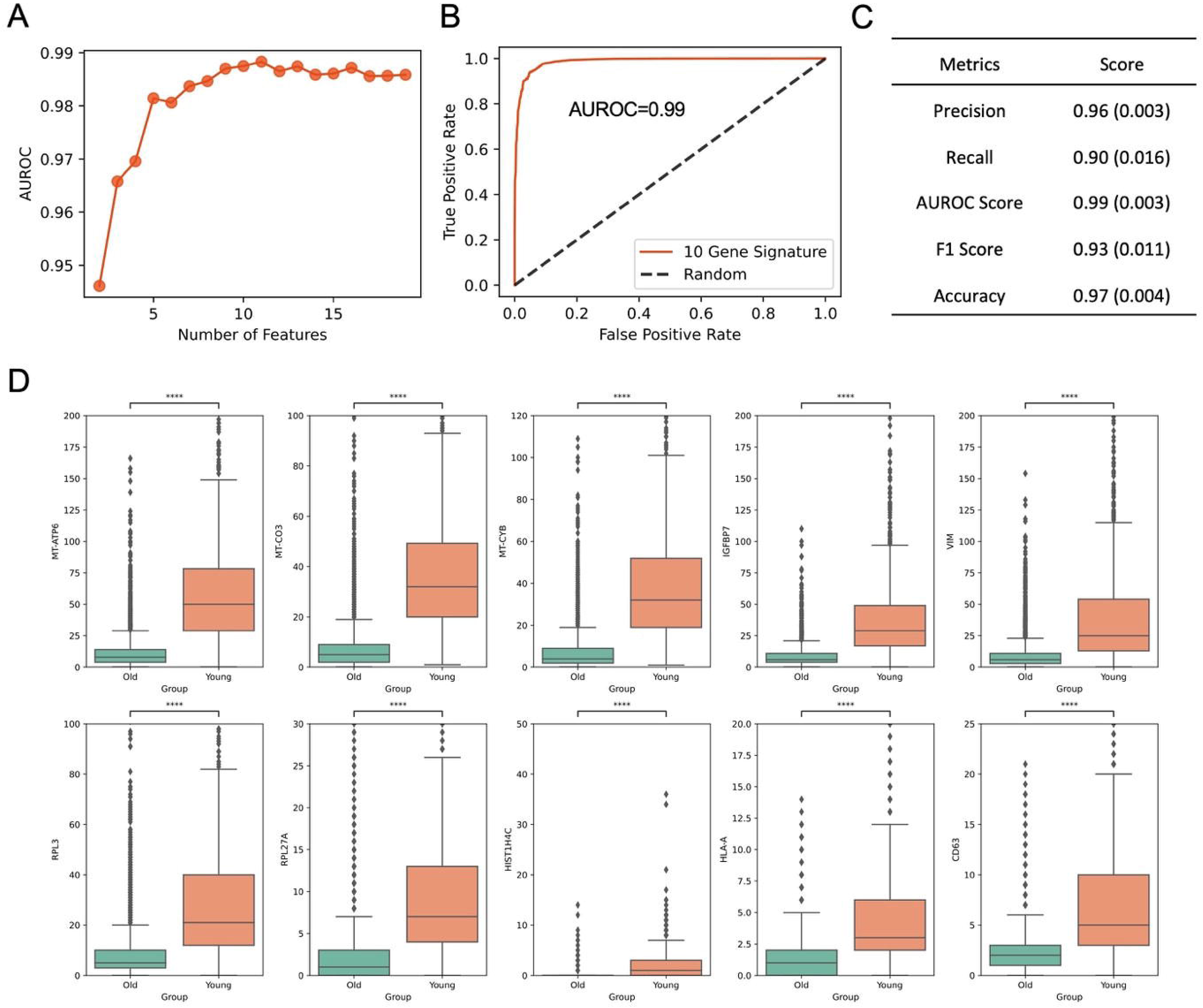
Old and young ovaries prediction based on a machine-learning algorithm. A. Cross-validated AUROC scores for different features included in the model. B. ROC curve for the predictive model using the 10-gene signature. C. Model evaluation metrics. Numbers outside (inside) the parentheses are the mean (standard deviation) of the 100 validation sets in 30 repeated runs of cross-validation. D. Expression of all genes in the signature between the old and young groups.

#### Profiling the cell-cell communication in young ovarian samples

To further investigate the mechanisms of ovarian aging, we examined the cross talk of various cell types. As mentioned earlier, we found that the Notch signaling pathway was significantly regulated in young oocyte cells and it also played critical roles in cell communications. Therefore, we used CellPhoneDB^28^ to profile the communication among cell types in the scRNA-seq data of young patients. The interactions among smooth muscle cells, endothelial cells and granulosa cells are the strongest, while others are relatively weaker (**Figure 6A, B**). Furthermore, we examined the interactions between oocyte and other cell types. Notably, several Notch family receptors were significantly enriched, which include Notch2, Notch3 and Notch 4 (**Figure 6C**). These pairs of interactions could play important roles in ovarian aging for oocytes to interacting with other cell types.

**Figure 6.**
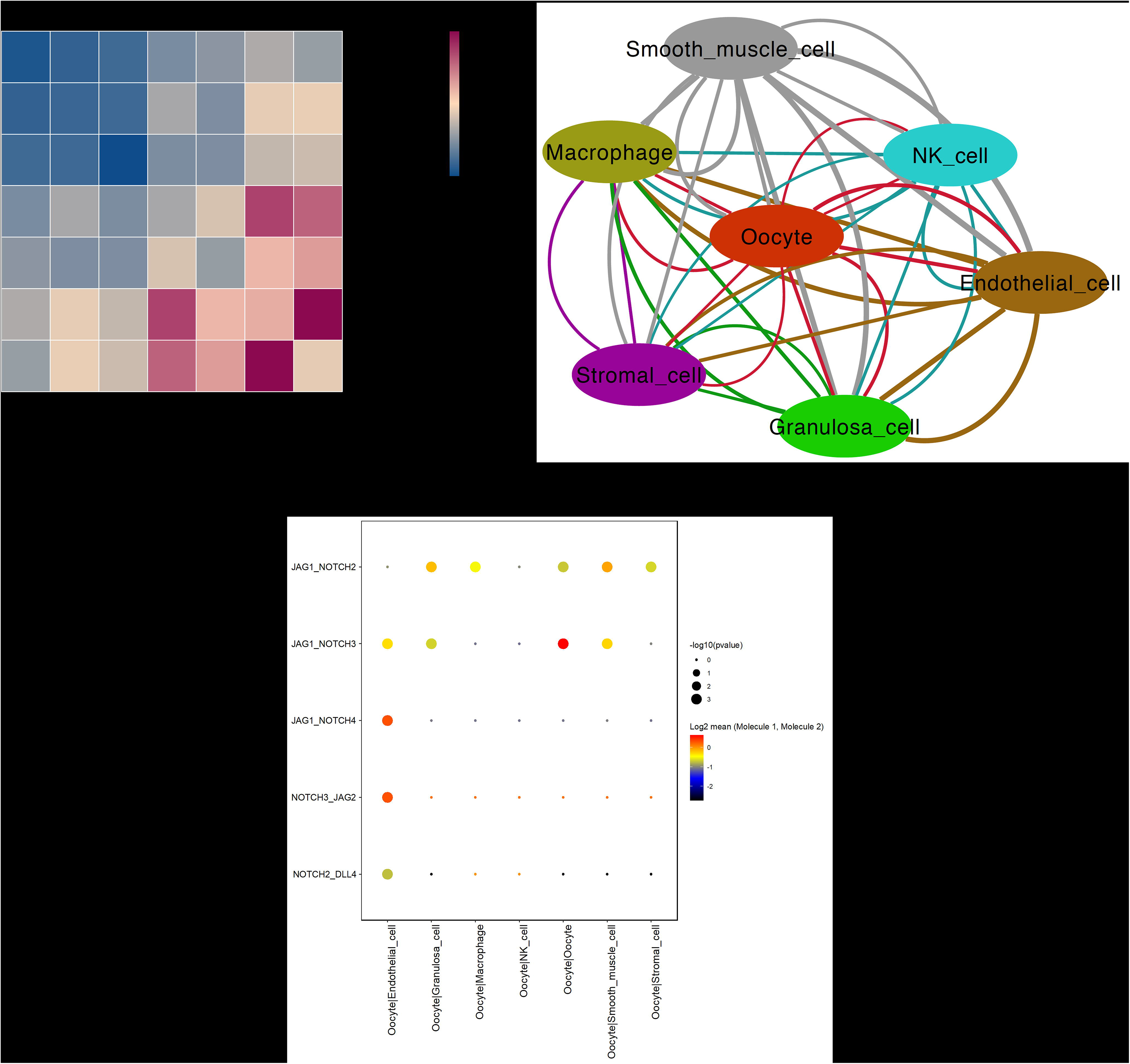
Cell interactions in young ovarian scRNA-seq data. A. Heatmap of interaction counts cell-cell interaction in young ovarian samples. B. Cell-cell interaction network in young ovarian samples. Colors and widths of edges represents number of interaction pairs between cell types. C. Bubble heatmap of Notch family receptors between oocyte and other cell types. Dot size represents - log10 p-value, while color represents log2 mean expression value of the receptor-ligand pair.

## Discussion

The ovary is characterized by highly heterogeneous populations at progressive developmental stages, and its cellular landscape is of indispensable importance to clarify the mechanism of ovarian aging. In this study, we integrated the scRNA-seq and scATAC-seq analyses to depict the transcriptional profiles and chromatin accessibility of old and young ovarian tissues. Unsupervised clustering of the scRNA-seq data revealed seven distinct cell populations. Annotation results were integrated with and confirmed by the scATAC-seq data. Marker genes, enriched pathways, and featured motifs for each cluster were identified. Notably, six subtypes of oocytes were recognized, representing different stages of oocyte development. Finally, a machine-learning algorithm was applied to calculate the optimal model for predicting ovarian aging.

The difficulty in procuring human ovary samples has long been an obstacle in revealing the cellular landscape of ovarian aging. Combining scRNA-seq and scATAC-seq data, we presented the first integrative study of ovarian aging at single-cell resolution. After multiple clustering and annotation, seven cell clusters were identified, including endothelial cells, granulosa cells, macrophages, NK cells, oocytes, smooth muscle cells, and stromal cells. This is consistent with the results of previous studies. Wang et al. performed the scRNA-seq analysis on the ovary samples collected from four young and four aged cynomolgus monkeys^7^. The same clustering results were found in their analysis, whereas the marker genes were slightly different^7^. Fan et al. applied scRNA-seq to human adult ovaries and depicted the molecular signature of growing and regressing cell populations^10^. Somatic cell types, including endothelial cells, granulosa cells, smooth muscle cells, and stromal cells, were also found in their study ^10^. In conclusion, the major cell types in human ovary samples were unveiled in this study.

The reclustering and cell trajectory analysis showed six subtypes of oocytes with distinct gene expression signatures during the sequential development stage. The differentially expressed genes were examined across the six subtypes for expression trends and oocyte development. Interestingly, TIMP1 and TIMP2 were negatively correlated with folliculogenesis. The TIMP gene family encodes proteins that inhibit matrix metalloproteinases (MMPs), thus preventing degradation of the extracellular matrix^29^. Previous studies have reported that TIMP1 and TIMP2 are involved in follicular development. Robinson et al. demonstrated that TIMP family members are localized in the oocyte cytoplasm and play an important role in the remodeling of the extracellular matrix during human gonadal development^30^. By investigating the gene expression levels of TIMP1 and TIMP2 in the granulosa and cumulus cells, Luddi et al. demonstrated that MMP and TIMP expression is involved in the regulation of oocyte maturation^31^. In this study, we showed that TIMP1 and TIMP2 were both significantly downregulated in the C4 and C5 oocytes (**Figure 2C and 2E**), which were mostly from old patients. This result indicates that the expression levels of TIMP1 and TIMP2 may determine the oocyte development stage as well as the aging process. Future studies should focus on deciphering the mechanistic roles of TIMP1 and TIMP2 in oocyte aging.

Chromatin accessibility analysis and robust peak identification revealed that the AP-1 TF family members, such as FOS, JUNB, FOSL1, and FOSL2 were highly enriched in the old oocyte cells compared to the young oocyte cells, consistent with the functional enrichment analysis and foot-printing analysis. Previous studies have reported that the AP-1 TFs participate in the early stages of murine follicle growth ^32^. Moreover, the AP-1 TF family has been reported to be involved in the aging of various cells. Shi et al. observed decreased AP-1 transcriptional activity in senescent human keratinocytes^33^. Medicherla et al. demonstrated that oxidative stress may contribute to impaired AP-1 binding activity, leading to adrenal aging in rats^34^. ATF3, an AP-1 TF, was found to be critical for the remodeling chromatin accessibility, thus promoting senescence of the human umbilical vein endothelial cells^35^. Herein, we demonstrated the crucial role of AP-1 genes in ovarian aging. Additionally, the Notch signaling pathway was found to be negatively regulated by the AP-1 complex ^36^. Chen et al. demonstrated that treatment with a Notch signaling pathway inhibitor led to a significant decrease in oocytes in the primordial follicles of newborn mice^37^, which implied that downregulation of the Notch signaling pathway is correlated with the dysfunction of the ovary. These studies indicated that the AP-1 TF family and its interaction with the Notch signaling pathway might regulate follicle formation and growth, thus contributing to ovarian aging. Further functional studies are needed to clarify their potential roles in the human ovary.

Finally, we built and optimized a predictive model based on the signature genes that were differentially expressed between the young and old ovaries. These genes include *MT-ATP6*, *MT-CO3, MT-CYB*, *IGFBP7*, *VIM*, *RPL3*, *RPL27A*, *HIST1H4C*, *HLA-A*, and *CD63*. The model presented excellent accuracy in cross-validation, suggesting its predictive and diagnostic significance for ovarian aging. Taken together, these observations provide novel insights into ovarian aging in humans. More detailed studies are essential to evaluate the function of the TIMP gene family and AP-1 TF family in oocyte development and maturation. Moreover, a predictive model based on machine learning promises to assist in the prediction of ovarian aging. Signature genes identified in this study may serve as new biomarkers and therapeutic targets for the diagnosis and treatment of age-related ovarian disorders.

## Material and methods

### Sample information

The experiments performed in this study were approved by the Ethics Committee of Affiliated Hospital of Nantong University. With informed consent, fresh ovarian tissues used for this study were from 6 donors who all underwent surgical treatment in Department of Obstetrics and Gynecology, Nantong University Affiliated Hospital between 2019-2020. All cases were excluded the use of hormones and malignant tumors. 4 cases menopausal women represent the elderly group (mean age 55.5 years, 50-62 years), who underwent whole uterus and double attachment resection because of uterine leiomyoma (n=3) and benign ovarian mass (n = 1). And the other 2 cases with normal fertility aged 31 and 34 years old who both underwent ovarian cyst removal surgery, we trimmed the irregular residual ovarian tissue to neat, and took a very small piece of normal ovarian tissue at the edge of the residual tissue during the operation. All the normal ovarian tissue were confirmed without histopathological abnormality by at least two independent pathologists.

### Single nuclei preparation and ATAC-library construction

Single nuclei ATAC-seq was performed following 10x Genomics NextGEM_SingleCell_ATAC_ReagentKits_v1.1 protocol. Homogenization was applied to tissues for breaking the tissues and obtaining the purified nuclei. Nuclei concentration was determined with hemocytometer under microscope and diluted in nuclei buffer (10x Genomics). The transposition experiments were then set up with thermocycler; The GEMs were then produced and cleaned up with the dynabeads, followed by performing the PCR reactions to add sample index. After quality control (QC) and quantification of the products, it was used for following library construction under the standard protocol (10x Genomics). The library was then sequenced using Illumina NovaSeq platform.

### Single cell preparation and library construction (for scRNA-seq)

Separated ovarian tissues were dissociated to single-cell resuspension for the single-cell transcriptome sequencing. The tissues were rinsed in cold DPBS for three times after dissection. Then the tissues were carefully transferred to digestion buffer (2mg/ml collagenase I, 1ml; 2mg/ml collagenase II, 1ml; 0.9U dispase, 0.5 ml; trypsin, 0.5 ml; DMEM, 1.5ml. Pre-heat to 37 degrees). The tissues were cut to appropriate smaller pieces in the digestion buffer and then gently shook in a 37-degree metal heater-shaker for 15 minutes. After the treatment, they were taken 10 μl from the resuspension and observed with a hemocytometer under microscope. If the cells were not sufficiently digested, it would be treated with digestion buffer for another 5 minutes. The cell suspension was then filtered by a 70-μm mesh filter, followed by washing the filter with another 5ml DMEM. The cells were concentrated by centrifugation (500g, 5 minutes at 4 degrees), and then the supernatant was removed. The cells were resuspended with 50 to 100 μl DMEM with 10% phosphate-buffered saline (PBS). The cells were then observed and counted with hemocytometer. The cell concentration was adjusted to 700 to 1200 cells per μl before loading to 10x Genomics Chromium machine.

The GEMs were produced and collected to perform reverse transcription for barcoding, and the first strand cDNA was purified with magnetic beads. After QC and quantification of the cDNA, it was then used for library construction under standard protocol (10X Genomics). The library was sequenced using Illumina NovaSeq platform.

### Single cell RNA-seq analysis

Seurat^38^ was used to read the Cellranger outputs (expression matrices) and used as the platform for downstream analysis. For quality control of each dataset, lowly-detected genes and cells with limited number of genes were discarded from the downstream analysis, in order to avoid the analysis driven by noise or low-quality cells. The gene filter was set to detection in at least 0.5% of all cells and the cell filter was set to a minimum number of 1000 of total expressed genes per cell. After the QC step, for each expression matrix, the expression value of each gene was normalized and transformed using default NormalizeData function in Seurat. Using the variation stabilizing transformation method (‘vst’ from Seurat), the top 2000 variable genes were selected in each matrix and were used as input for the ‘FindIntegrationAnchors’ function of Seurat. The expression matrices were then integrated with the ‘IntegrateData’ function. The integrated data were dimension reduced with principal component analysis (PCA; top20 dimensions). In the PCA space, nearest neighbors were defined among cells with KNN method (‘FindNeighbors’, top 15 PCs were selected) and cells were then grouped with Louvain algorithm (‘FindClusters’ in Seurat, resolution equal to 0.5). UMAP dimension reduction was then performed using ‘RunUMAP’ function for visualization purpose. CellPhoneDB was used to profile the communication among cell types in the scRNA-seq data of young and old patients^28^.

### Detection of differentially expressed genes (DEGs)

DEGs were first computed to test significantly highly expressed genes for each cluster. For each cluster, only the genes expressed in more than 25% of that group were considered. All other cells were used as the background. For statistical test, we used the ‘MAST’ method implemented in Seurat. DEGs were defined as genes whose log fold-change were over 0.4 compared to the background, and with a q-value (FDR) smaller than 0.05.

In each cell type, differentially expressed genes of the cells from tissues with primary tumors and those with recurrent tumors were computed. The method is the same as above described except that log fold-change threshold were set to 0.25.

### Single-cell ATAC-seq analysis

Raw scATAC-seq data was first processed by the ‘cellranger-atac’ tool downloaded from 10x Genomics, which mapped the sequenced reads to the hg38 genome reference and generated the fragment files. R package ‘archR’ was then used for the downstream analysis with fragment files as input^20^. For quality control, cells with less than 4000 unique fragments or TSS enrichment scores less than 8 were removed. Doublets were also removed by using the default settings. Cells passing the filters were then clustered by using the ‘Seurat’ method implemented in the ‘archR’ package with high resolution values (1.5 or 2). UMAP with 30 neighbors was used for dimension reduction and visualization. For cell cluster annotation, we first integrated the cell annotations generated by previous scRNA-seq data by using the ‘addGeneIntegrationMatrix’ function of ‘archR’ and then further curated the cell cluster annotations by specific marker genes. For peak calling, we used the recommended ‘MACS2’ with default settings on aggregated coverage profiles of identified cell populations. For the rest downstream analysis of motif enrichment, footprinting analysis and trajectory plots, we used the functions provided by ‘archR’ packages with the recommended settings.

### Statistical Analysis and machine learning based prediction model

Mutual information scores were used to evaluate feature importance. We also considered permutation importance score to select features, which were outperformed by mutual information in model accuracy benchmarking. Each evaluation score was averaged over 100 repeated runs. Statistical models including random forest classifier, gradient boost classifier, XGB classifier, Naïve Bayes were considered. We used a genetic algorithm (TPOT) to select the best model and optimize its hyperparameters. The area under the curve of the receiver operating characteristic curve (AUROC), was used to evaluate model performance. We also calculated other metrics for comparison purposes. All models were evaluated using 5-fold cross validation with stratified train-test splits that preserve the percentage of samples for the prediction target. All metrics were averaged over 30 repeated runs.

### Hematoxylin-eosin staining and Immunofluorescence

Histologic analysis was performed on ovarian tissues using hematoxylin and eosin staining (H&E). After operation, ovarian tissues were fixed in 4% formaldehyde and processed for routine paraffin embedding after fixation. We took 5-mm-thick sections for hematoxylin and eosin staining. Tissue sections were deparaffinized and rehydrated in graded ethanol solutions. In order to repair the antigen, the sections were immersed in antigen retrieval solution (P0088, Beyotime Biotechnology) and heated in 97 °C water bath for 20 minutes. First of all, immunol staining blocking buffer (0.5% triton X-100, 5% Horse serum and PBS) was used before staining, which can reduce non-specific staining. Immunostaining was performed using the primary antibody against c-Fos (1:300, Abcam, ab208942, Cambridge, USA) and c-Jun (1:300, Abcam, ab40766, Cambridge, USA) at 4°C overnight. On completion of the incubation at 4°C, tissues were washed three times with PBS. Then localization of c-Fos was monitored using Alexa Fluor 594 (Goat Anti-Mouse, 1:500, Abcam, ab150116, Cambridge, USA) and Alexa Fluor 488 (Goat Anti-Rabbit, 1:500, Abcam, ab150077, Cambridge, USA) for c-Jun. After 2 hours of co-cultivation at room temperature, tissues were washed 3-5 times with PBS for 10 minutes each time. Finally, antifade mounting medium with DAPI (P0131, Beyotime Biotechnology) was used to stain the nucleus. The tissues were observed under Zeiss microscope and the images were processed by ZEN software.

## Supporting information

H&E staining for the collected tissue.

Functional enrichment analysis of marker genes for each cell type.

Quality control of scATAC-seq analysis.

Doublets removal by ArchR for four old patient samples

Doublets removal by ArchR for two young patient samples.

Browser track showing the scATAC-seq signal of FOSL2 and ATF3 across old and young patient samples.

## Acknowledgement

We appreciate the assistance received from Intanx Life (Shanghai) Co. Ltd. for data processing and consultation.

---

Supplementary Figure 1: H&E staining for the collected tissue.

Supplementary Figure 2: Functional enrichment analysis of marker genes for each cell type.

Supplementary Figure 3. Quality control of scATAC-seq analysis.

A. scATAC-seq signals of the unique fragment number per cell for old and young groups.

B. scATAC-seq signals of old and young groups are highly enriched on TSS.

C. Fragment size distribution for old and young groups.

Supplementary Figure 4. Doublets removal by ArchR for four old patient samples.

Supplementary Figure 5. Doublets removal by ArchR for two young patient samples.

Supplementary Figure 6. Browser track showing the scATAC-seq signal of FOSL2 and ATF3 across old and young patient samples.

