## Supplementary figures and images for "Integrated analyses of the single-cell ATAC-seq and RNA-seq reveal the epigenetic landscape of human ovarian aging"

### Browser track showing the scATAC-seq signal of FOSL2 and ATF3 across old and young patient samples.

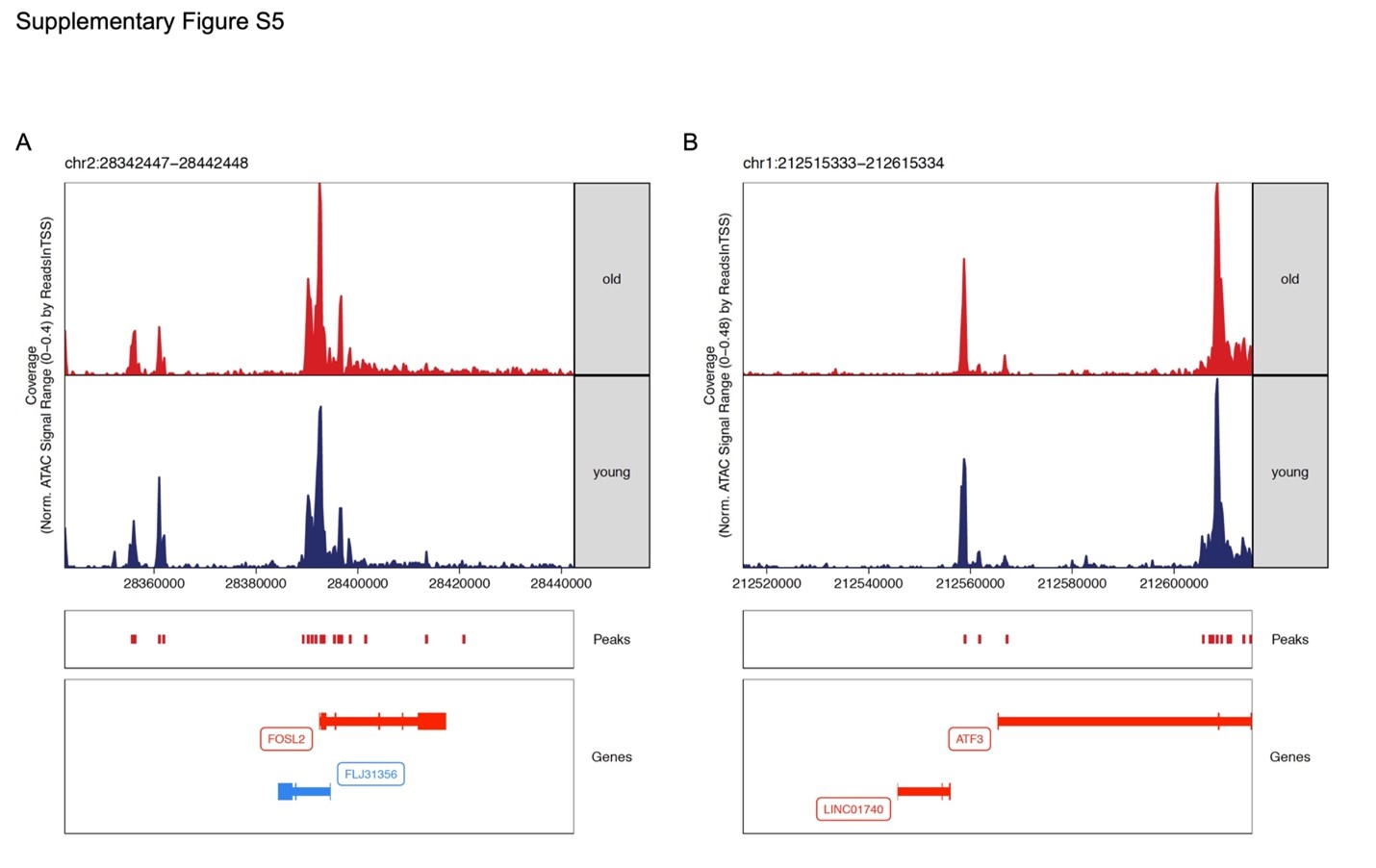

### Doublets removal by ArchR for four old patient samples

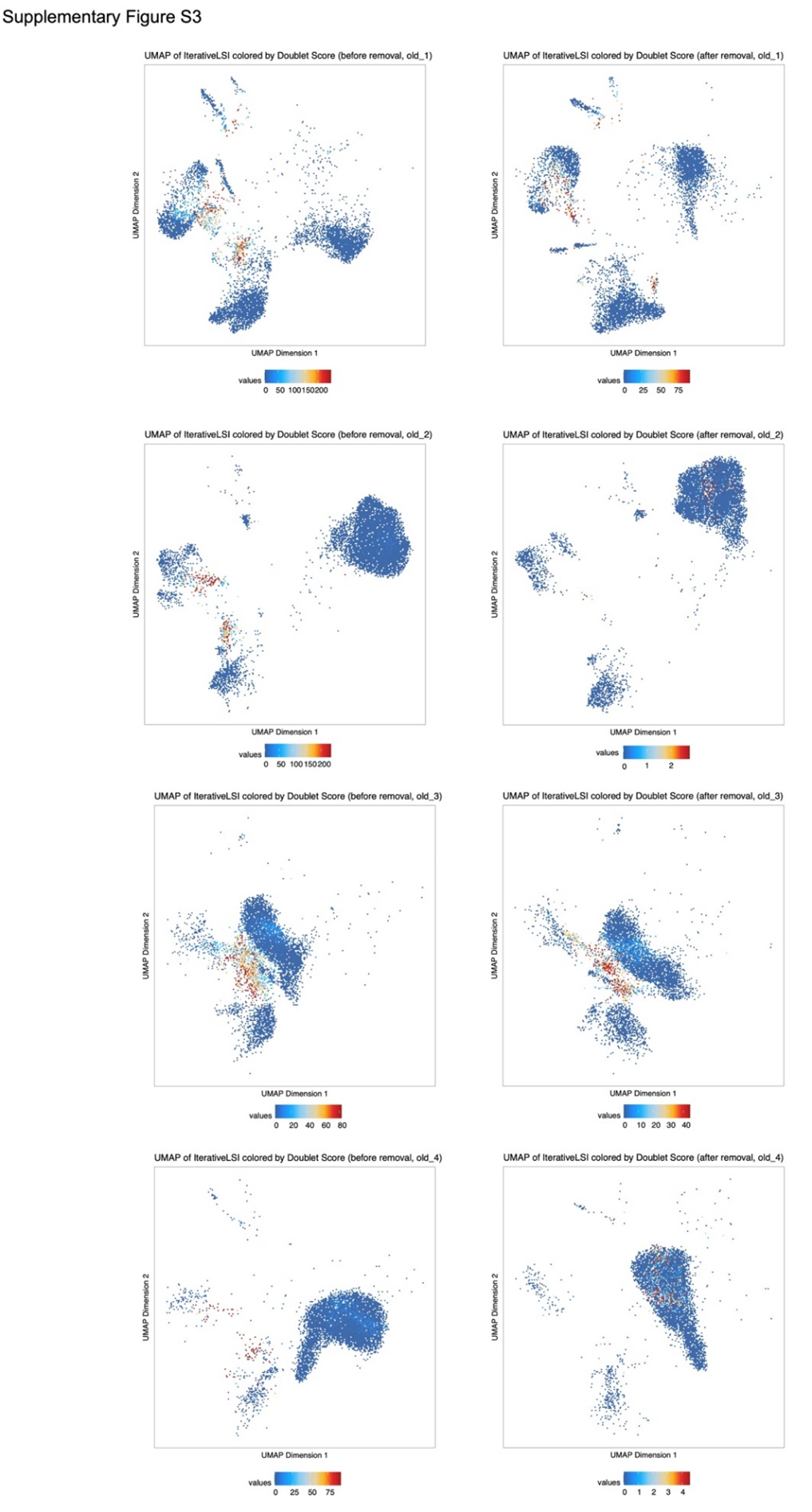

### Doublets removal by ArchR for two young patient samples.

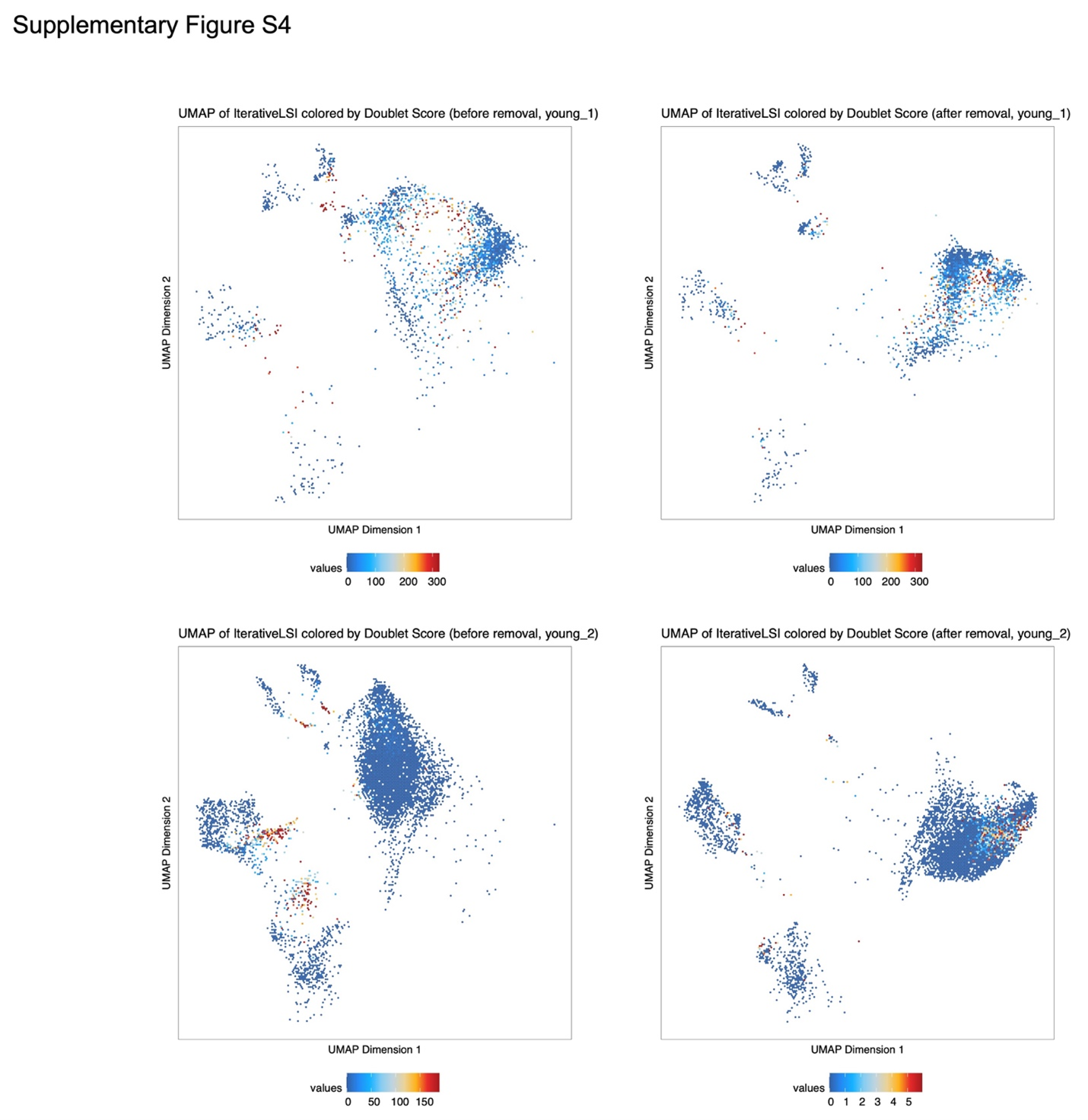

### Functional enrichment analysis of marker genes for each cell type.

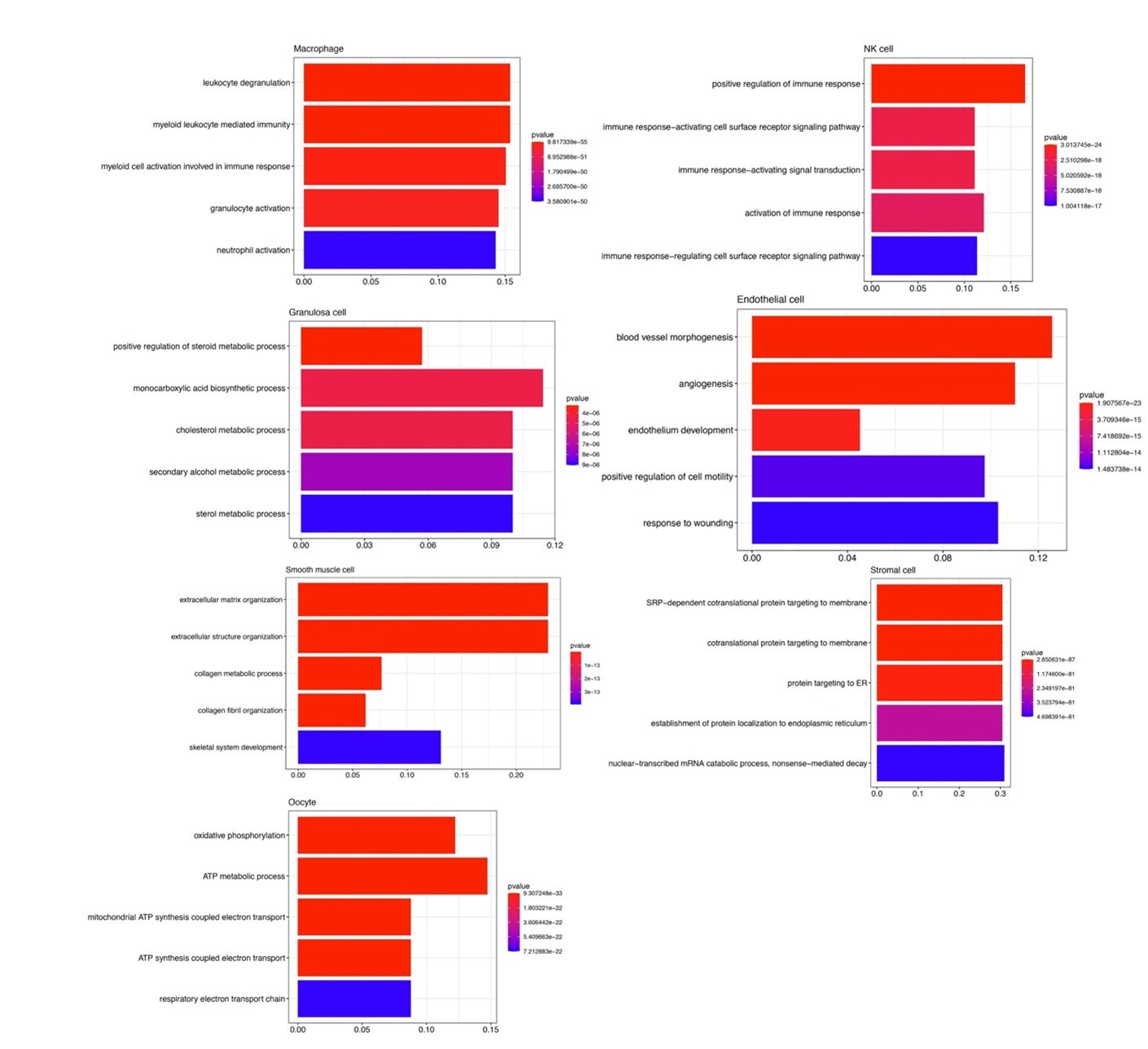

### H&E staining for the collected tissue.

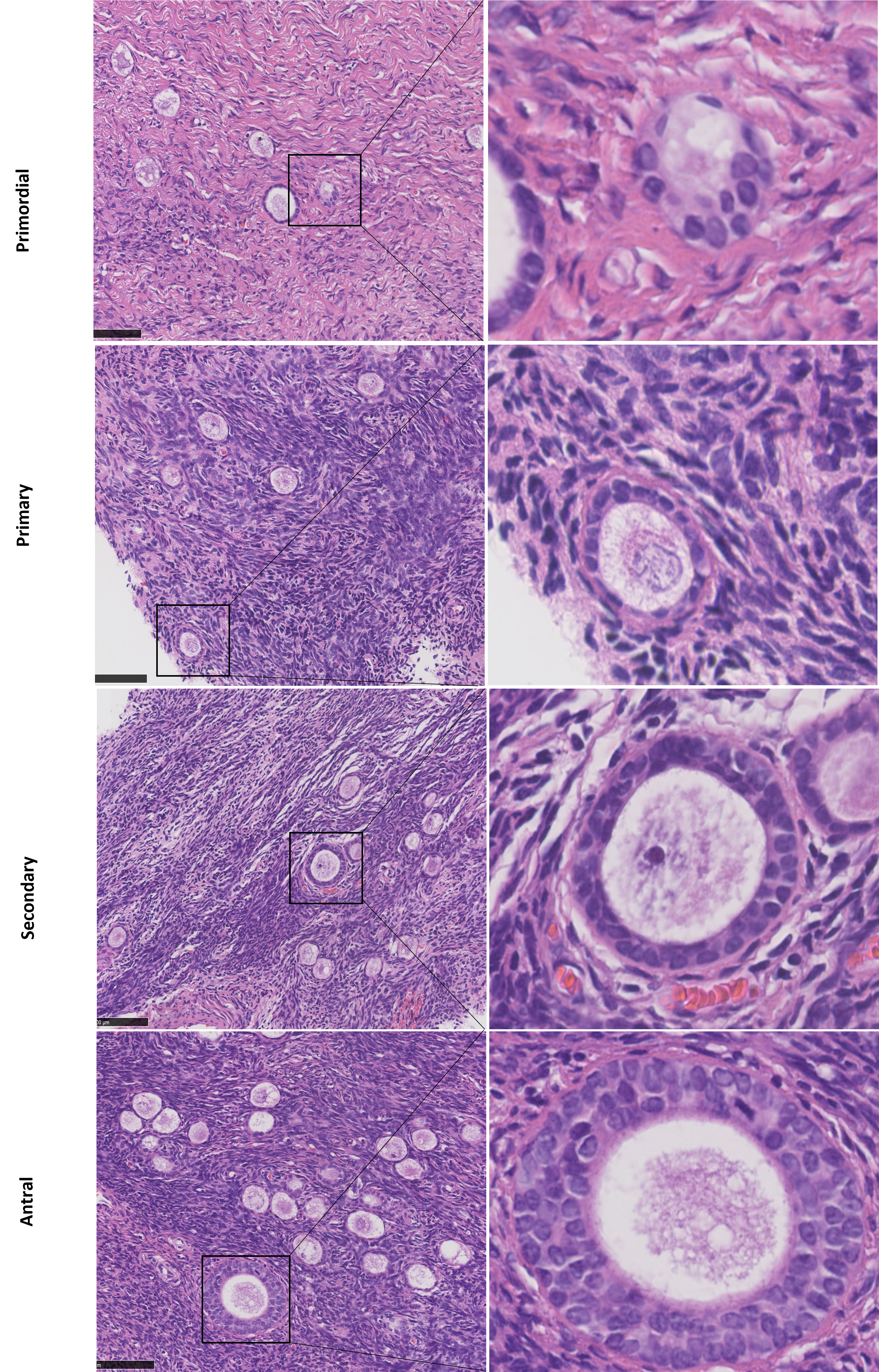

### Quality control of scATAC-seq analysis.

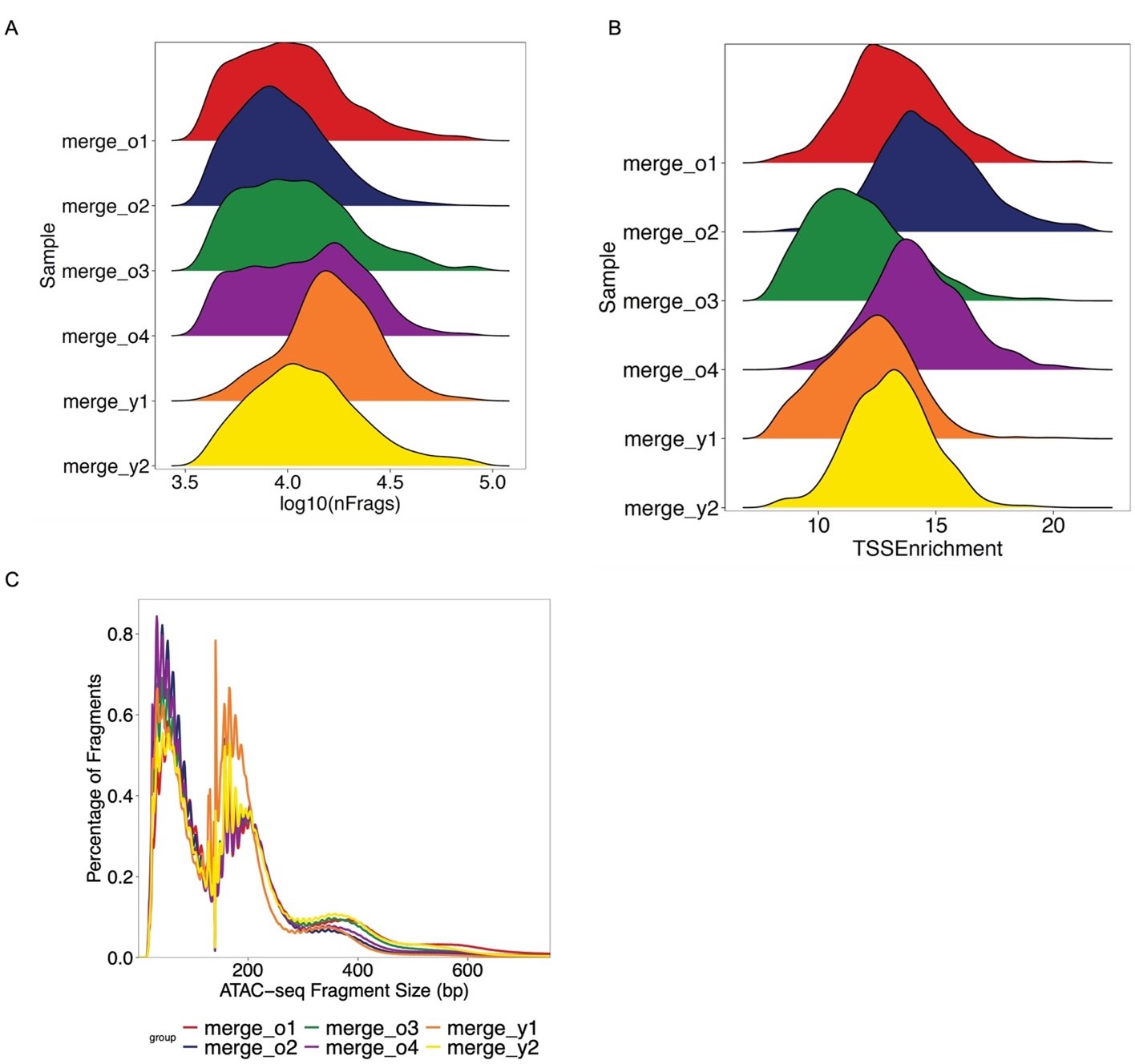
